# Non-destructive tree volume estimation using mobile laser scanning: Impact of the tree shape on measurement error

**DOI:** 10.64898/2026.08.11.744158

**Authors:** Justin Holvoet, Philippe Lejeune, Jérôme Perin, Bastien Vandendaele, Gauthier Ligot

**Affiliations:** Liège University, Faculty of Gembloux Agro-Bio Tech, Passage des Déportés, 2, 5030, Gembloux, Belgium; Natural Resources Canada, Canadian Forest Service, Laurentian Forestry Centre; 1055 Rue du Peps, Québec, QC G1V 4C7, Canada

**Author notes:** Corresponding author. Email adress.

## Abstract

Accurate tree volume estimation is central to forest management and carbon accounting. Allometric equations are widely used but fall short when used out of their intended scope, regions and environmental conditions. Mobile laser scanning offers an alternative to estimate volume based on direct measurement of tree geometry, but how tree shape affects its measurement error remains poorly documented. We compared stem diameters, total tree height and merchantable stem volume derived from mobile laser scanning with destructive measurements on 176 felled trees of eight species (four hardwood and four softwood) in 32 plots in Wallonia, Belgium. Trees were scanned with a Zeb Horizon RT scanner. Volume was estimated with a quantitative structure model and with a taper equation fitted to the point cloud diameters. Height was compared with Vertex IV clinometer measurements. The impact of tree shape descriptors derived from the point clouds on the errors were where tested with mixed-effects models that account for the nesting of measurements within trees, plots and species.

Across 7,218 diameter measurements, the mean error was +0.5 cm (−0.3 cm for hardwood, +1.3 cm for softwood), and precision declined above 15 m. For hardwood, mobile laser scanning gave more precise heights than the clinometer (relative root mean square error of 6.9% vs. 8.8%), but it was slightly less precise for softwood (7.4% vs. 6.1%). Volumes from the quantitative structure model were underestimated by 21% on average, and a correction factor removed most of this bias but requires reference data to calibrate. The taper-based volumes were nearly unbiased overall, with a relative root mean square error of 15.7%. Processing took between 1h15 and 2h35 per plot and about seven minutes per tree.

Tree shape variables explained only 9–23% of the variance in errors (29–39% when species and plot were included). Errors were larger within the crown, and volume and height errors were linked to crown size and shape, and to tree size. Tree shape was not identified as the main source of measurement uncertainty. Future work should test scanning protocol, stand conditions and processing steps together with tree characteristics to allow large-scale use in forest inventories.

## 1. Introduction

Accurate assessment of individual tree wood volume is fundamental for environmental, economic, and forest management purposes (Mura et al., 2025). Reliable volume estimation enables optimal forest resource utilization and effective monitoring of forest dynamics. Additionally, systematic errors in volume estimation can lead to economic losses and unsustainable management (Sladek and Neruda, 2007).

In commercial forestry, merchantable wood volume is a key determinant of timber market value, making accurate measurements critical for forest owners, logging companies, and timber buyers. Reliable volume estimates help reduce conflicts between suppliers and customers, support transparent transactions, and improve forest inventory consistency (Mura et al., 2025). From a management perspective, correct estimation of standing tree volume supports sustainable forest planning by informing harvest schedules, thinning operations, and reforestation strategies. Overestimation can result in overharvesting and depletion of forest resources, and may constrain economic returns and biased growth and yield projections (Fan et al., 2020).

Tree volume measurement also plays a pivotal role in environmental monitoring and carbon accounting, as it is a prerequisite to estimate aboveground biomass and carbon storage (Gschwantner et al., 2019; Longuetaud et al., 2013). Accurate volume estimates are crucial for quantifying forest carbon stocks, improving climate models, and ensuring the credibility of carbon credit schemes. Errors in volume estimation can propagate into significant biases in carbon sequestration assessments, ultimately affecting environmental reporting and climate policy (Zhou et al., 2022). Consequently, robust and accurate volume and biomass estimation remain central to sustainable forest management, carbon stock assessment, and climate change mitigation efforts (Hossain, 2025).

In this situation, allometric equations are often the chosen solution to estimate wood volume on large samples of trees over a large territory. Despite their widespread use, gaps in the development and application of allometric equations continue to limit the accuracy of forest resource assessments and carbon stock estimations (Mahmood et al., 2016). A persistent issue is their use outside of their intended scope. In many cases, practitioners must rely on equations developed for other species, forest types, or geographic regions, which increases the uncertainty and potential bias in volume and biomass estimates (Mulatu et al., 2024; Yang et al., 2023). Many classical allometric equations were derived from relatively small and geographically restricted datasets, leading to poor representation across tree species, age classes, and diameter ranges. Applying such models beyond their original calibration domain often introduces systematic bias and reduces predictive reliability (Sebrala et al., 2022).

Empirical evidence also suggests that several established volume models exhibit structural inconsistencies. For example, volume equations used in Finland over the past four decades, although generally robust, displayed illogical behavior for small trees, particularly for models relying solely on diameter at breast height (DBH) and total stem height (Kangas et al., 2022, 2020, Laasasenaho, 1982). Similarly, the polynomial models produced by Dagnelie and used in Belgium have shown reduced accuracy for large trees (Servotte, 2017).

Recent studies indicate that tree stem form can evolve over time and will vary spatially due to environmental and management-related factors. Kangas et al., (2020) reported that, for the major tree species in Finland, newer datasets revealed taller and more slender trees than older datasets. They attributed this shift primarily to denser stands and changing silvicultural practices. However, regional differences suggest that climatic and environmental conditions may also influence allometric relationships. Thus, we can suspect that recent and future climate change will impact trees allometry increasing the need for updated allometric equation and direct measurement tools such as 3D scanners.

In this field of research related to forest inventory, Terrestrial laser Scanning (TLS) have been widely used to estimate individual stem merchantable volume, which correspond to the volume of the stem from the base up to the point of the stem which reach a circumference of 20 cm (Calders et al., 2020). TLS is a ground-based laser scanning technique in which a scanner, mounted on a tripod at fixed positions, produces a dense and highly accurate 3D point cloud of the surrounding objects. By combining scans acquired from several positions within a plot, TLS allows a detailed, non-destructive reconstruction of stem form and volume. Due to the logistical constraints and difficulties to integrate TLS in an operational forest inventory, Mobile laser scanning (MLS) is often preferred, even if less precise, as it makes data acquisition easier to perform and decreases the occlusion in the scan (Holvoet et al., 2025a). MLS, on the contrary of TLS, is able to continuously record the surroundings while the operator walks through the plot, producing a 3D point cloud much faster than TLS, though with somewhat lower precision.

Based on these two technologies, stem volume can be derived from a point cloud in several ways. One approach relies on quantitative structure models (QSM), which reconstruct the structure of a tree with a set of cylinders fitted to the point cloud (Raumonen et al., 2013). The merchantable stem volume is then obtained by summing the volumes of the cylinders along the stem, up to the point where the circumference falls below 20 cm. QSMs were initially developed and validated on TLS data. A second approach is based on stem taper. Diameters are measured at successive heights along the stem on the point cloud, a taper function is fitted to these diameters, and the function is integrated along the stem to obtain the volume. This approach relies directly on measured diameters rather than on a full reconstruction of the tree, and it makes it possible to limit the influence of aberrant measurements, which are frequent in the upper part of the stem (Pitkänen et al., 2021).

In their review, (Holvoet et al., 2025a) reported relative RMSE% for the MLS ranging from 8.9% to 21.82% and bias ranging from 0.6% to 14.55% when compared with destructive stem volume measurements. Stovall et al., (2023) used a Zeb1 (GeoSLAM®) MLS to estimate stem volume of 20 felled trees pertaining to the following species: *Acer rubrum*, *Pinus strobus*, *Quercus rubra* and *Tsuga canadensii*. Using DBH-derived stem taper, they obtained accurate volume estimates up to 11.5 m above ground, although accuracy decreased progressively with height along the stem. They also reported that softwood species were more difficult to measure because dense needles reduced stem visibility and limited laser penetration into the crown. Vandendaele et al., (2024) used a Hovermap (Emesent Pty Ltd) scanner in a Canadian hardwood forest in leaf off conditions to measure 26 trees pertaining to the species *Acer saccharum, Abies balsamea* and *Betula alleghaniensis*. They noted that MLS tend to overestimate the total merchantable volume of smaller trees while it underestimated larger trees. They also noted that stem volume was consistently underestimated when using quantitative structure model (QSM) to estimate the volume. Chiappini et al., (2022) compared the volume of 50 felled pine trees in a pine dominated forest with MLS estimations and found a bias of 4.1% and RMSE% of 12,4% using the RANSAC method (De Conto et al., 2017). The results obtained in these studies suggest that MLS volume estimation can produce more accurate estimates than commonly used non-destructive approaches.

While volume measurement from TLS and MLS point clouds has been repeatedly validated against destructive reference measurements (Calders et al., 2015; Demol et al., 2021; Stovall et al., 2023; Vandendaele et al., 2022), a systematic evaluation of how tree-level characteristics specifically affect MLS measurement accuracy remains absent from the literature. Most validation studies have reported overall accuracy metrics at the tree or plot level, without assessing the contribution of species type, tree shape, and architecture to the observed errors. A study by Holvoet et al., (2025b) suggested that MLS measurement accuracy may vary between softwood and hardwood species, particularly for stem volume, but also possibly for structural attributes such as crown dimensions or total tree height. However, quantifying these differences rigorously requires destructive reference measurements across a large set of trees covering a broad range of species, sizes, and structural forms. Beyond species group alone, tree-level characteristics such as crown and branching complexity, stem convexity or stem to crown ratio may influence MLS measurement errors, and their relative contribution has yet to be assessed.

This study aims to i) evaluate the accuracy and precision of the MLS against destructive reference measurements for: Diameters measured along the stem, total tree height, and stem merchantable volume and ii) determine which tree-level characteristics influence the accuracy of these measurements.

## 2. Material and method

### 2.1 Study site

A total of 32 circular plots (18 m radius) were established in mixed hardwood and softwood forests across Wallonia, Belgium (Figure 1). Each plot was surveyed using a Zeb Horizon RT MLS (FARO®, Lake Mary, FL, USA) to acquire three-dimensional point clouds of the forest structure

**Figure 1:**
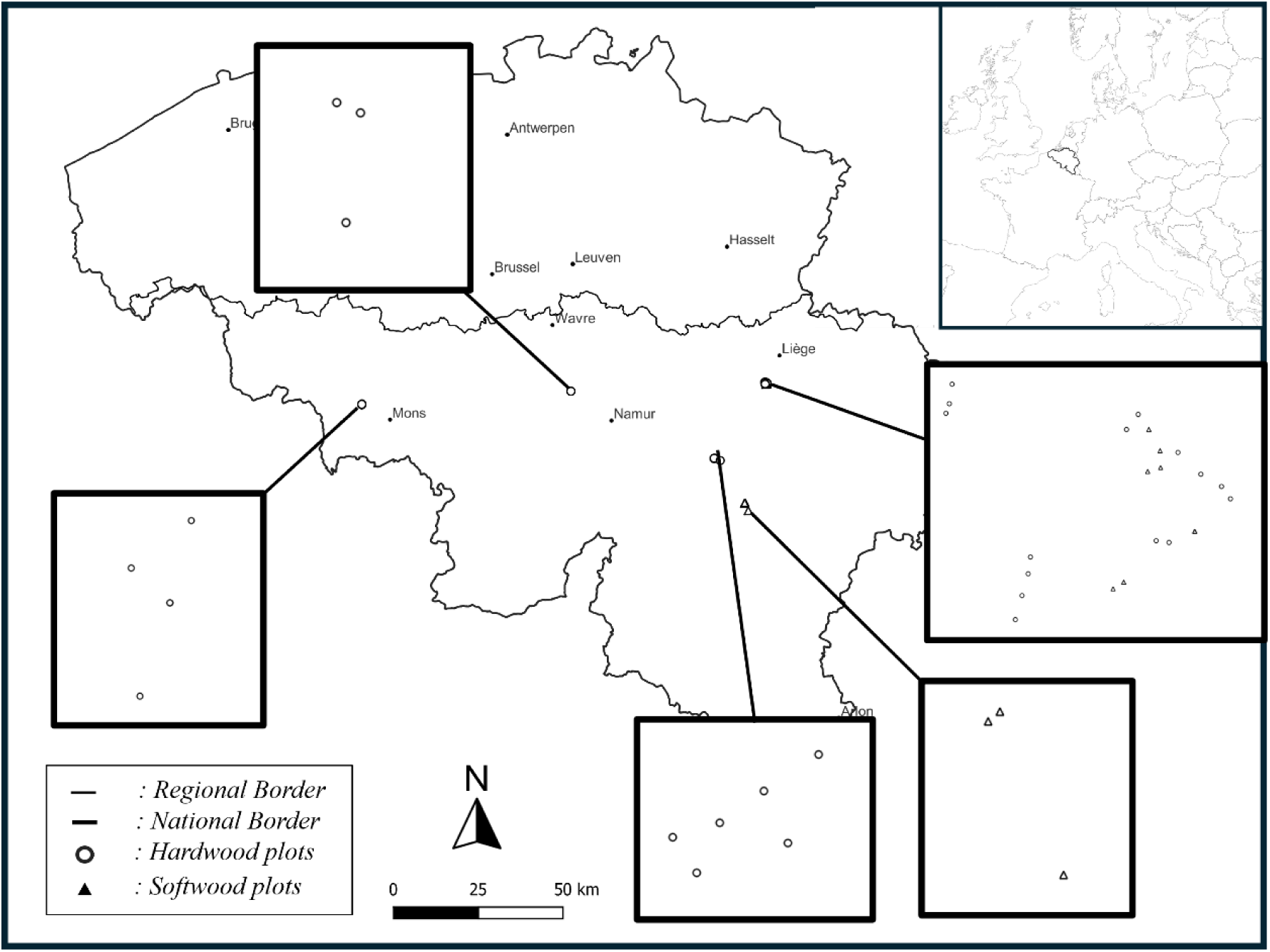
Location of the 32 circular plots (18 m radius) where trees were scanned with mobile laser scanning and then felled, in Wallonia, Belgium.

Within these plots, 176 trees were selected and logged. Prior to harvesting, tree height was measured in the field using a Vertex IV electronic hypsometer (Haglöf Sweden AB, Sweden). Following the scanning campaign, all selected trees were felled. Detailed measurements, including stem volume by successive diameter measurement and total tree height, were collected manually on the ground and used as reference data for the evaluation of point cloud-derived measurements.

The number of sampled trees varied substantially among species, ranging from 7 individuals for cypress to 37 for beech. Tree diameter at breast height (DBH) classes ranged from 10 to 80 cm. However, both the distribution of individuals across DBH classes and the evenness of these distributions differed markedly among species. Some species were represented in only two DBH classes, whereas others spanned nearly the entire range. Across all species, most sampled trees fell within the 10 to 40 cm DBH classes (Table 1).

**Table 1:**
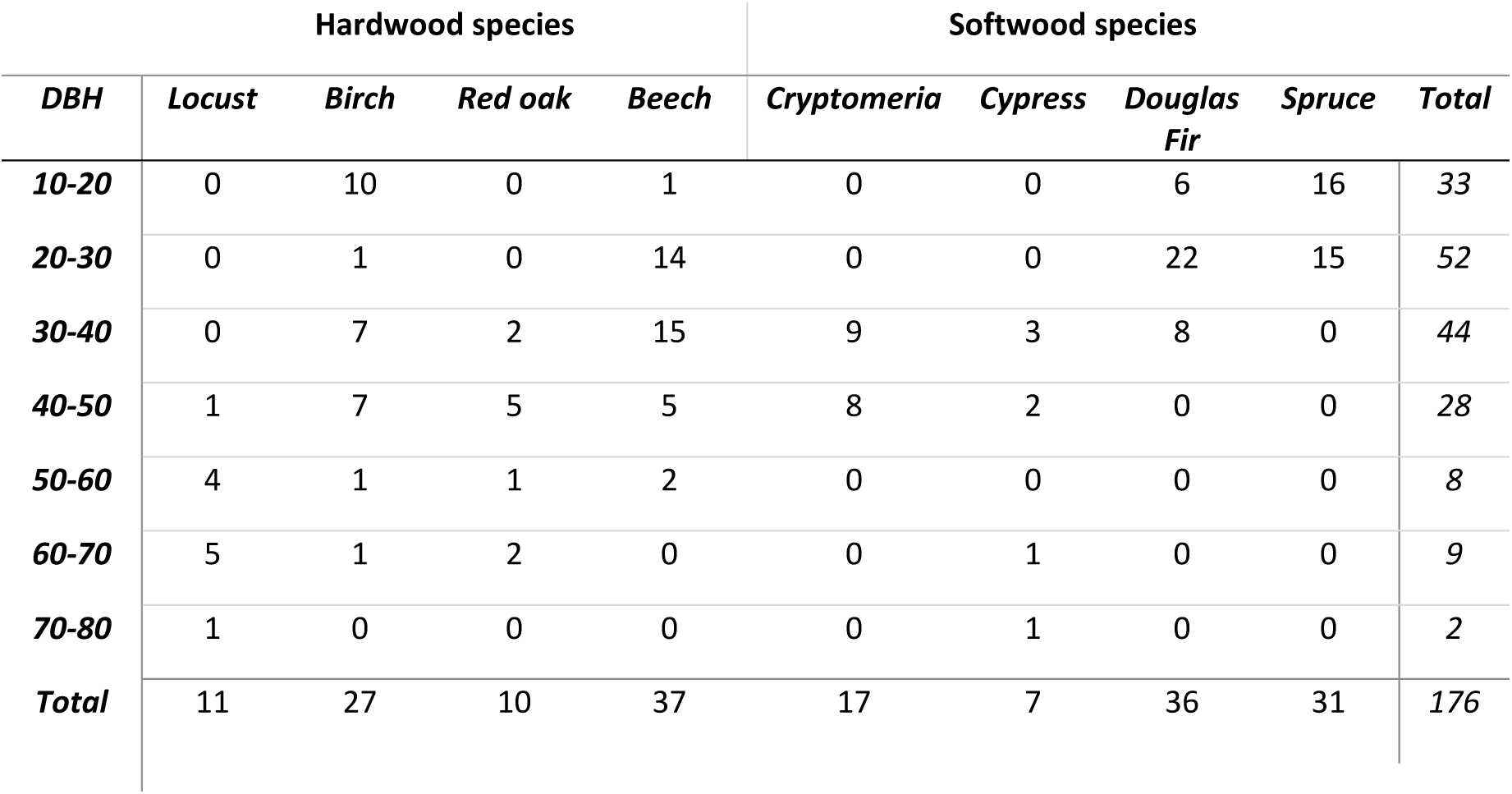
Distribution of the scanned and felled trees in diameter at breast height (DBH) classes.

### 2.2 Destructive measurements

Measurements were conducted on all 176 sampled felled trees. For each tree, stem length, stem merchantable volume, and the number of large branches were recorded. Before felling, the relative position of each tree in the plot was measured using azimuth and distance.

After felling, reference tree height was defined as the rectilinear distance between the base of the stem and the apical bud. This distance was measured using a Vertex electronic clinometer, and the height of the remaining standing stump was added to obtain the total stem length.

Large branches were identified as any branches with a circumference greater than 20 cm at their insertion point on the stem, whether living or not. The number of large branches and their position along the stem were recorded for each tree.

Stem circumference was measured successively along the stem. Circumference was measured with a tape at 1-m intervals from the stem base up to the point where the circumference dropped under 30 cm. Beyond this point, measurements were taken every 30 cm until the stem circumference decreased under 20 cm after which measurement were stopped. Where circumference was unobtainable due to the contact of the stem with the ground, the diameter was measured in place using a caliper. In branching sections, the stem was defined as the largest of the two resulting sections.

Stem merchantable volume was defined as the volume of the stem from the base of the tree up to the circumference of 20 cm. To calculate it, the volume of each stem section was first calculated using the Smalian formula.

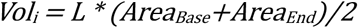

*Where L is the length of the measured section, Area_Base_ is the Area of the base of the section and Area_End_ is the Area of the end of the section*

The total stem merchantable volume was then obtained by summing up the volume all stem sections.

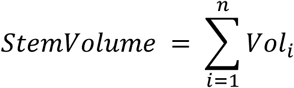

### 2.3 MLS scans

The scan were conducted using a Zeb Horizon Rt Scanner (Faro). The scanner properties can be found in table 2.

**Table 2:** Specifications of the ZEB Horizon RT mobile laser scanner (FARO) as given by the manufacturer. SLAM: simultaneous localisation and mapping; mrad: milliradian.

| Scanner model | ZEB HORIZON RT |
| --- | --- |
| Range (m) | 100 |
| Beam divergence (mrad) | 3.0 |
| Accuracy (cm) | 3 |
| Measurement rate (point/sec) | 300,000 |
| Wavelength (nm) | 903 |
| SLAM algorithm | GEOSLAM |

MLS data were acquired on 18-m-radius plot in January-March 2025. The scans were carried out following the methodology presented in Holvoet et al (2025b) (see figure A1). Before scanning, 16 metal poles were placed respectively at 15 and 22m at North, West, South, East, North-West, South-West, North-East, and South-East. Additionally, a white sphere was placed at the center of the plot and another 5m north from the center in order to facilitate the navigation of the operator in the plot. Prior scan setup took around 15 minutes while scanning operations took around 15 to 25 minutes per plot.

#### Point cloud processing

Tree-level point cloud measurements included total tree height and stem volume. In addition, several metrics describing tree architecture and crown structure were derived from the individual tree point clouds, as detailed in Table 4. MLS measured trees were matched with manual measurements based on stem X/Y position and diameter as described in Holvoet et al., (2025b).

The point clouds were segmented into individual tree point clouds using the Simple Forest plugin (Hackenberg et al., 2021) on the Computree platform (Othmani et al., 2011).

After denoising, SimpleForest builds a digital terrain model (DTM) from the ground points, and normalize point heights. To identify potential stem candidates, a thin horizontal slice of the point cloud was extracted at breast height above the DTM. Within this slice, points belonging to stems were retained and clustered, so that each valid cluster serves as a stem seed for one tree. The remaining vegetation points were then assigned to these clusters to create individual tree point clouds.

To remove the noise from the point clouds, a statistical outlier removal filter (SOR) was applied. SOR, when applied to whole tree point clouds at once, tend to over-filter smaller stems due to the heterogeneity in point density (Charlebois et al., 2026). To avoid this issue, we decided to subdivide the individual tree point clouds in 3m high slices and to then apply the SOR filter to each slice with the following parameters (k=50, m=0.5) (Fig A2)

**Figure 2:**
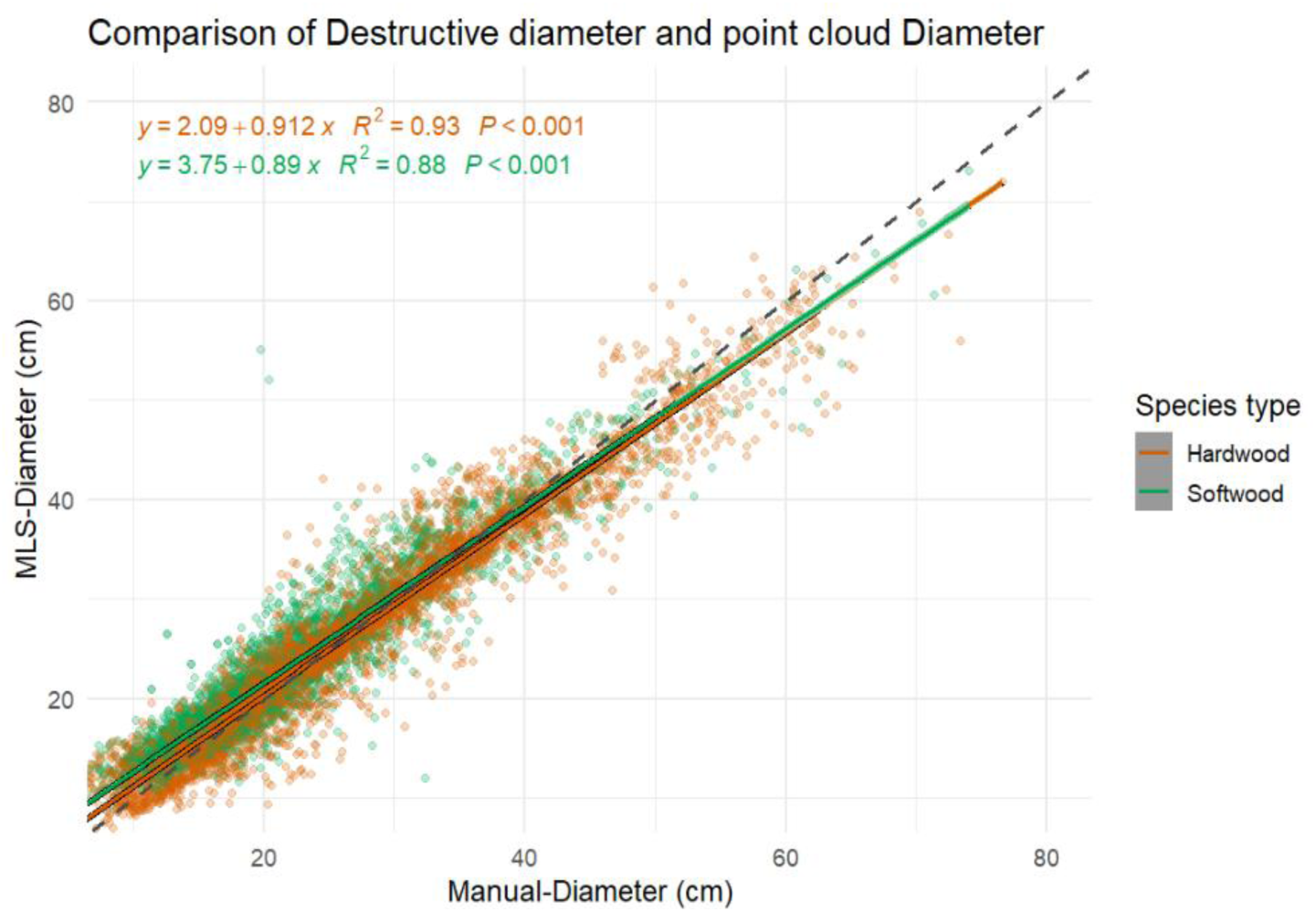
Comparison of the diameters estimated from mobile laser scanning point clouds (MLS-Diameter) against diameters measured on the felled trees (Manual-Diameter), for hardwood and softwood, based on 7,218 measurements from 176 trees. The dashed line is the 1:1 line and the solid lines are linear regressions per species group. MLS : Mobile Laser Sanning

Total tree height was estimated as the difference between the minimum and maximum z-coordinates of the individual tree point cloud. Point cloud-derived tree heights were compared with the tree height estimates measured on standing trees using a Vertex IV electronic hypsometer and with the reference measurements carried out on felled trees.

Stem merchantable volume was estimated using two different methods. The first method consisted of the construction of a QSM using TreeQSM version 2.3.2 (Raumonen et al., 2013) with the following input parameters : PatchDiam1 = 2, PatchDiam2Min = 3, PatchDiam2Max = 2. Using QSM, the merchantable volume corresponded to the cumulative volume of all stem cylinders up to the first cylinder to go under 20 cm in circumference. The height of the remaining stump on the field was removed from the first cylinder.

The second method segmented the stem points from the individual tree point clouds using the QSM as reference for stem position. The points present within 10 cm distance of each stem cylinder were kept and labelled as stem points. Successive diameters of the stem point cloud were then calculated using the ITSMe R package on 10 cm large slices at 30 cm intervals (Terryn et al., 2023). Aberrant diameter values were defined as value exceeding the previous measurement by 30% as defined by Kangas et al., (2020) in case of multiple aberrant measurements in a row, the last validated measurement was used as reference. Additionally, another measurement of 0 cm of circumference was added at the total tree height previously measured. This additional measurement allowed to make up for the impossibility to measure accurate diameter close to the top of the stem. This additional measurement helped the fitted taper model to converge better toward the diameter of 0 around the total tree height.

Based on the point cloud diameter measurements, the following model was fitted, and each parameter estimated for each tree to obtain individual tree taper:

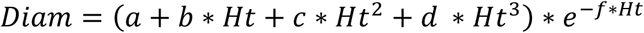

*Where Diam is the estimated diameter of the stem at a given height, Ht is the height at which the diameter is estimated, and parameters a,b,c,d,f are constants fitted for each individual tree, and e is the Euler number*.

The model was not selected to represent a biologically correct stem taper, but it was empirically built to fit to the observed diameters while decreasing the weight of extreme and aberrant measurements. The parameters were estimated separately for each tree by non-linear least squares, using the Levenberg–Marquardt algorithm implemented in the *nlsLM* function of the *minpack.lm* R package (Elzhov et al., 2023). The fitting used the diameters measured on the 10 cm point cloud slices (after removal of aberrant values) and the additional zero diameter placed at the total tree height.

The fitted taper equation was subsequently used with the following formula to obtain stem volume:

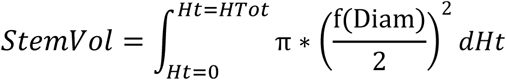

*Where StemVol is the stem volume, Ht is the height of the diameter estimation, f(Diam) is the fitted model, and HTot is the total tree height*.

Measurement errors were calculated as the MLS or Vertex IV estimate minus the destructive reference measurement: diameter error (DiamError), MLS and Vertex IV height errors (MLS_HtError and Vertex_HtError), and taper-based stem volume error (VolError). Positive values therefore indicate overestimation.

The average processing time for every step is displayed in Table 3

**Table 3:** Time taken for the processing steps. SLAM : simultaneous localisation and mapping

| **Step** | **Scale** | **Processing time** |
| --- | --- | --- |
| **Plot setup** | Plot | 10-15 minutes |
| **Scan** | Plot | 15-20 minutes |
| **SLAM algorithm** | Plot | ~30 to 60 minutes |
| **Plot point cloud segmentation** | Plot | 30~60 minutes |
| **Total for the plot** | Plot | 1h15-2h35 |
| **Cleaning the individual point cloud** | Tree | ~30 seconds |
| **Building the QSM** | Tree | ~5 minutes |
| **Shape parameters measurement** | Tree | ~1 minute |
| **Diameter, height and volume measurements** | Tree | 30 seconds |
| **Total for a tree** | Tree | ~7 minutes |

### 2.4 Analysis of the effect of tree shape on measurement errors

#### Exploratory analysis

The dataset has a nested structure: the 7,218 diameter measurements were taken along the stems of 176 trees, which were located in 32 plots and belonged to eight species. Observations are not entirely independent, since measurements from the same tree, plot or species are likely to share common sources of error. The correlation analyses described below ignore this structure and were computed on pooled observations (7,218 diameters or 176 trees). They are used only to explore the data. Their coefficients should be read as descriptive and not as formal tests. On the contrary, the mixed effect models produced to predict the error of measurement based on the shape parameter could take this nested structure into account by including plot ID, tree species and tree ID as random factors.

Several additional metrics describing tree architecture and crown structure were derived from the individual tree point clouds. These variables are listed and described in Table 4. A more thorough description of all variables is available in table A1. Relationships between tree structural variables and measurement errors were first explored with Pearson correlation coefficients computed between all variables (Table 4) and the errors, using the *cor()* function of R (R Core Team). The matrices were displayed as heat maps. In addition, principal component analyses (PCA) were run with the FactoMineR package (Lê et al., 2008) on standardized variables (mean 0, standard deviation 1), separately for the diameter errors (7,218 measurements) and for the height and volume errors (176 trees). These PCA are presented in Appendix A.

**Table 4:** Tree shape parameters used to explain measurement errors. ITSMe: Individual Tree Structural Metrics R package (Terryn et al., 2023); MLS: mobile laser scanning.

| **Variable** | **Abbreviation** | **Description** |
| --- | --- | --- |
| **Species type** | SpTp | Hardwood or softwood |
| **Number of large branches** | NbBr | Number of branches with a circumference above 20 cm at insertion |
| **Circumference at breast height** | CBH | Stem circumference at 1.30 m (cm) |
| **Height of diameter measurement** | MsHeight | Height above the stem base at which a diameter was measured (m) |
| **Measurement section** | MeasSect | Diameter measured on the stem or within the crown |
| **Stem convexity** | Convexity | Ratio between the area of the spline fitted on the stem slice at breast height and the area of its convex hull (Bauwens et al., 2021) |
| **Stem complexity** | Complexity | Ratio between the circumference of the circle and the perimeter of the spline fitted at breast height |
| **Crown projected area** | CPA | Area of the convex hull of the crown points projected on a horizontal plane (m<sup>2</sup>) |
| **Crown volume** | CrVol | Volume of the alpha shape around the crown points (m<sup>3</sup>) |
| **Crown proportion** | CrownProp | Proportion of the total tree height occupied by the crown |
| **Crown point density** | CrPtDensity | Number of crown points divided by crown volume (points m<sup>-3</sup>) |
| **Crown shape** | CrShape | Crown volume divided by crown projected area (m) |

#### Mixed-effect models

Linear mixed-effects models were fitted to four response variables: the diameter error (DiamError), the Vertex IV height error (Vertex_HtError), the MLS height error (MLS_HtError) and the taper-based stem volume error (VolError). For each response, the candidate fixed effects were the species type (hardwood or softwood) and the tree architecture variables listed in Table 4 (CPA, CrVol, CrShape, CrownProp, CBH, NbBr, Convexity, Complexity and CrPtDensity). For the diameter model, the height of the measurement (MsHeight, tested as a second-order polynomial) and the measurement section (MeasSect: stem or crown) were also included. A measurement was classified as a crown measurement when it was located above the crown base height estimated with the ITSMe R package (Terryn et al., 2023). All continuous predictors were centered and scaled to a mean of 0 and a standard deviation of 1, so that coefficients are comparable across predictors.

Plot identity and species were tested as random factors in all models. For the diameter model, tree identity (nested within species) was also tested, because several diameters were measured along each stem. Models were fitted with the lme4 package (Bates et al., 2015) in R, and p-values of fixed effects were obtained with the lmerTest package (Kuznetsova et al., 2017) using Satterthwaite’s approximation. Starting from the full model, non-significant fixed and random terms were removed one at a time, starting with the least significant, until all remaining fixed effects were significant with a 95% interval. The explanatory power of each final model was assessed with the marginal R^2^ (variance explained by fixed effects) and the conditional R^2^ (variance explained by fixed and random effects), following Nakagawa and Schielzeth (2013) and Nakagawa et al. (2017). The intraclass correlation coefficient (ICC) quantified the share of variance attributable to the random effects.

## 3. Results

### 3.1 Accuracy of MLS estimates against reference destructive measurements

#### Diameters

A total of 7,218 stem diameter measurements were available for comparison between the manual measurements and the one made on the 10cm slices of point cloud (Table 5). Overall, diameter estimation exhibited low bias (0.5 cm) although standard deviation increased substantially along the stem (Figures 2 and 3). MLS-diameters are slightly overestimated for softwood and slightly underestimated for hardwood.

**Figure 3:**
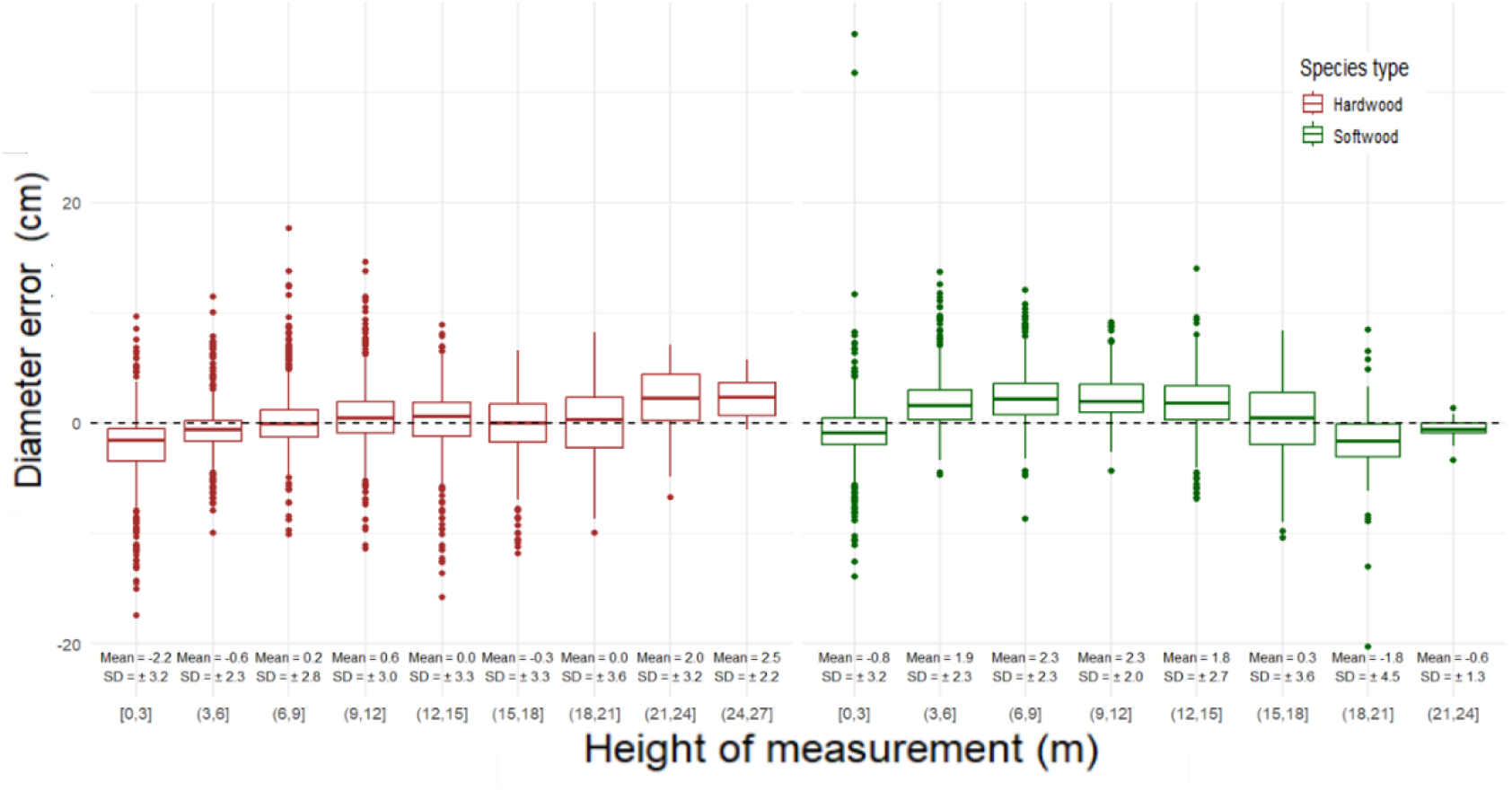
Diameter error along the stem for hardwood (left) and softwood (right) trees, by class of measurement height. The error is estimated as the diameter estimated from the mobile laser scanning point clouds minus the diameter measured on the felled trees, so positive values indicate overestimations. Boxes show the median and the first and third quartiles. The mean and standard deviation for each height class is indicated at the bottom of the figures.

**Table 5:** Mean stem diameter error, mean relative error and standard deviation for all species, hardwood species and softwood species. Diameters were measured up to the diameter of 7 cm. Values are the mean with their S5% confidence interval in parentheses

|  | **All Species** | **Hardwood** | **Softwood** |
| --- | --- | --- | --- |
| **Number of observations** | 7218 | 3787 | 3431 |
| **Mean diameter error (cm)** | 0.5 (0.4 – 0.5) | -0.3 (-0.4 – -0.2) | 1.3 (1.2 – 1.4) |
| **Mean relative error (%)** | 4.40 (4.05 – 4.76) | 0.71 ( 0.248 – 1.17) | 8.34 (7.83 – 8.86) |
| **Standard deviation of the diameter difference (cm)** | 3.18 | 3.16 | 2.99 |

Diameter measurements were underestimated within the first 3 m above ground (mean bias of −2.2 for hardwood, and –0.8 for softwood cm) and overestimated between 6 and 15 m (figure 2). Above 3 m, the bias remained stable; however, measurement precision progressively decreased with increasing stem height, with a marked deterioration observed above 15 m with standard deviation increasing above 3.3 cm for hardwood and 3.6 cm for softwood. This trend is more pronounced for softwood than for hardwood, where measurements are more stable (figure 3)

#### Height and volume

Across all species confounded, both methods exhibited low mean relative errors and comparable overall accuracy (Table 6 and Figure 4). However, a slight systematic bias was observed: the Vertex IV measurements slightly overestimated tree height (mean error = 0.17 m; p.value = 0.000225; mean relative error = 0.43%), whereas MLS-Height estimates were slightly underestimated (mean error = −0.54 m; p.value = 0.223; mean relative error = −2.34%). Despite these differences in bias direction, both approaches yielded similar overall precision, with %RMSE values of 7.60% for Vertex IV and 7.13% for MLS.

**Figure 4:**
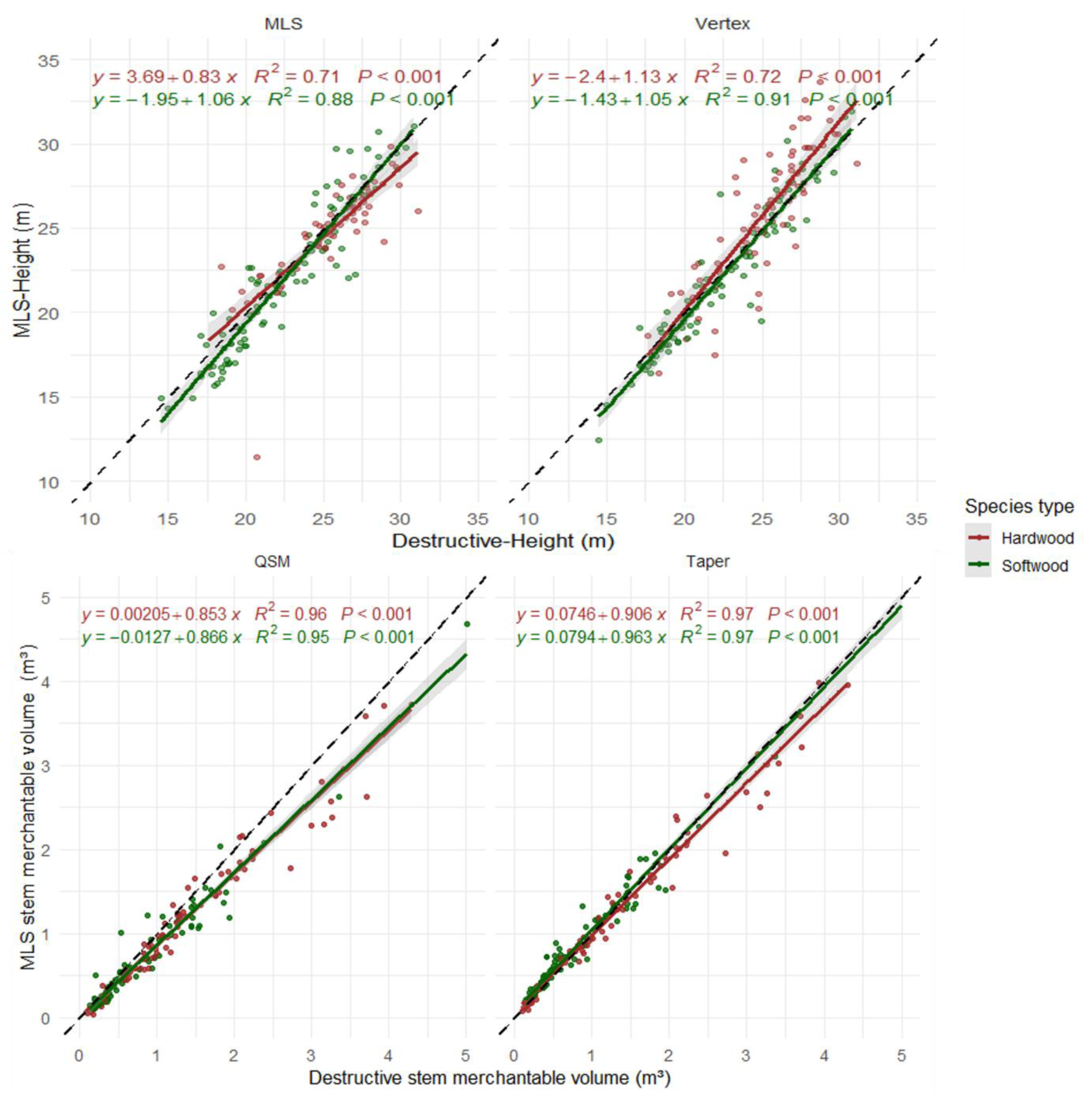
Comparison of estimated and reference values. (a) Total tree height estimated with mobile laser scanning, (b) total tree height measured with the Vertex IV clinometer, (c) merchantable stem volume estimated with a quantitative structure model (QSM), and (d) merchantable stem volume estimated by integrating the taper model fitted to the point cloud diameters. Reference values are the destructive measurements on the 176 felled trees. The dashed line is the 1:1 line; points are colored by species group. MLS: Mobile laser scanning, QSM: Quantitative structure model.

**Table 6:** Accuracy and precision of total tree height estimated with the Vertex IV clinometer and with mobile laser scanning, for all species, hardwood and softwood, relative to the height measured on felled trees. The error is the estimate minus the reference. Values are the means with their 95% confidence interval in parentheses. RMSE%: root mean square error as a percentage of the mean reference height.

|  | **Vertex** | **Point cloud** |
| --- | --- | --- |
| **Mean height error for all species (m)** | 0.17 (-0.11 – 0.45) | -0.54 (-0.79 – -0.30) |
| **Mean relative error (%)** | 0.43 (-0.72 – 1.57) | -2.34 (-3.45 – -1.26) |
| **Standard deviation of the height error for all species (m)** | 1.79 | 1.60 |
| **RMSE% for all species (%)** | 7.60 | 7.13 |
| **Mean height error for hardwood species (m)** | 0.75 (0.27 – 1.24) | -0.59 (-0.98 – -0.21) |
| **Mean relative error for hardwood species (%)** | 2.87 (0.90 – 4.83) | -2.15 (-3.84 – -0.46) |
| **Standard deviation of the height difference for hardwood species (m)** | 2.09 | 1.64 |
| **RMSE% for hardwood species (%)** | 8.78 | 6.88 |
| **Mean height error for softwood species (m)** | -0.30 (-0.58 – -0.02) | -0.50 (-0.83 – -0.18) |
| **Mean relative error for softwood species (%)** | -1.54 (-2.76 – -0.31) | -2.49 (-3.97 – -1.06) |
| **Standard deviation of the height error for softwood species (m)** | 1.35 | 1.58 |
| **RMSE% for softwood species (%)** | 6.14 | 7.36 |

When separating species groups, contrasting patterns emerged between hardwood and softwood stands. For hardwood species, Vertex IV overestimated tree height (0.75 m; 2.87%), while MLS underestimated it (−0.59 m; −2.15%). Among hardwood species, MLS provided better performance in terms of %RMSE (6.88%) compared to Vertex IV (8.78%), indicating higher precision.

In contrast, for softwood species, both methods consistently underestimated tree height (bias of −0.30 m for Vertex IV and −0.50 m for MLS). Here, Vertex IV performed slightly better in terms of %RMSE (6.14%) compared to MLS (7.36%).

The performance of the two stem volume estimation approaches, QSM, and the point cloud-based stem taper method (Taper-Volume), is summarized in Table 7, while the relationship between estimated and reference stem volumes is illustrated in Figure 4.

**Table 7:** Accuracy and precision of merchantable stem volume estimated with a quantitative structure model (QSM), and the taper-based method, for all species, hardwood and softwood, relative to the volume measured on felled trees. The error is the estimate minus the reference. Values are the means with their 95% confidence interval in parentheses. RMSE%: root mean square error as a percentage of the mean reference volume.

|  | **QSM** | **Stem taper** |
| --- | --- | --- |
| **Mean Volume difference for all species ($\text{m}^3$)** | -0.207 ( -0.241 – -0.171) | -0.001 (-0.026 – 0.024) |
| **Mean relative error (%)** | -21.2 (-24.0 – -18.3) | 6.50 (3.7 – 9.3) |
| **Standard deviation of the volume difference for all species ($\text{m}^3$)** | 0.234 | 0.168 |
| **RMSE% for all species (%)** | 29.1 | 15.7 |
| **Mean volume difference for hardwood species ($\text{m}^3$)** | -0.262 (-0.317 – -0.207) | -0.057 (-0.096 – -0.017) |
| **Mean relative error for hardwood species (%)** | -22.0 ( -25.3 – -18.7) | -2.8 (-5.3 – -0.3) |
| **Standard deviation of the volume difference for hardwood species ($\text{m}^3$)** | 0.255 | 0.183 |
| **RMSE% for hardwood species (%)** | 26.2 | 13.7 |
| **Mean volume difference for softwood species ($\text{m}^3$)** | -0.156 (-0.196 – -0.113) | 0.048 (0.022 – 0.079) |
| **Mean relative error for softwood species (%)** | -20.5 (-24.9 – -15.8) | 15.1 (10.9 – 19.3) |
| **Standard deviation of the volume difference for softwood species ($\text{m}^3$)** | 0.202 | 0.136 |
| **RMSE% for softwood species (%)** | 32.9 | 18.7 |

The QSM approach systematically underestimated stem volume with the underestimation increasing approximately linearly with stem volume (figure 4). Consequently, for all species combined, mean QSM-Volume error was −0.207 m^3^ while the mean relative error was −21.2%, resulting in the highest overall error among the two evaluated methods (%RMSE = 29.1%).

The taper-volume method provided accurate stem volume estimates (Table 7), with a mean volume error of −0.000994 m^3^ across all species. Although this method exhibited a slight positive mean relative error (6.50%), it achieved the lowest overall %RMSE (15.72%) and the lowest standard deviation of volume errors (0.168 m^3^), indicating the highest overall precision and consistency. Species-specific analyses revealed contrasting performances between hardwood and softwood trees. For hardwood species, the taper-volume method produced accurate estimates, with the lowest %RMSE (13.7%) and the smallest variability in volume error (0.183 m^3^), despite a slight negative bias (mean relative error = −2.82%).

For softwood species, the taper-volume yielded a lower %RMSE (18.7%) although it exhibited a larger positive bias, with a mean volume difference of 0.0477 m^3^ and a mean relative error of 15.1%.

### 3.2 Exploratory analysis of the impact of tree shape on measurement errors

#### Diameters

The exploratory analysis showed that the structural variables were organized along gradients of tree size, crown architecture, stem surface complexity and measurement height, but diameter errors were only weakly associated with them (PCA, figure A3). The correlation analysis confirmed that no structural variable was strongly related to DiamError (all |r| < 0.22; Figure 5). The strongest associations were with CrownProp, MsHeight, CrPtDensity and CBH, which suggests that diameter errors are not driven by a single tree-level attribute but by several weak factors.

**Figure 5:**
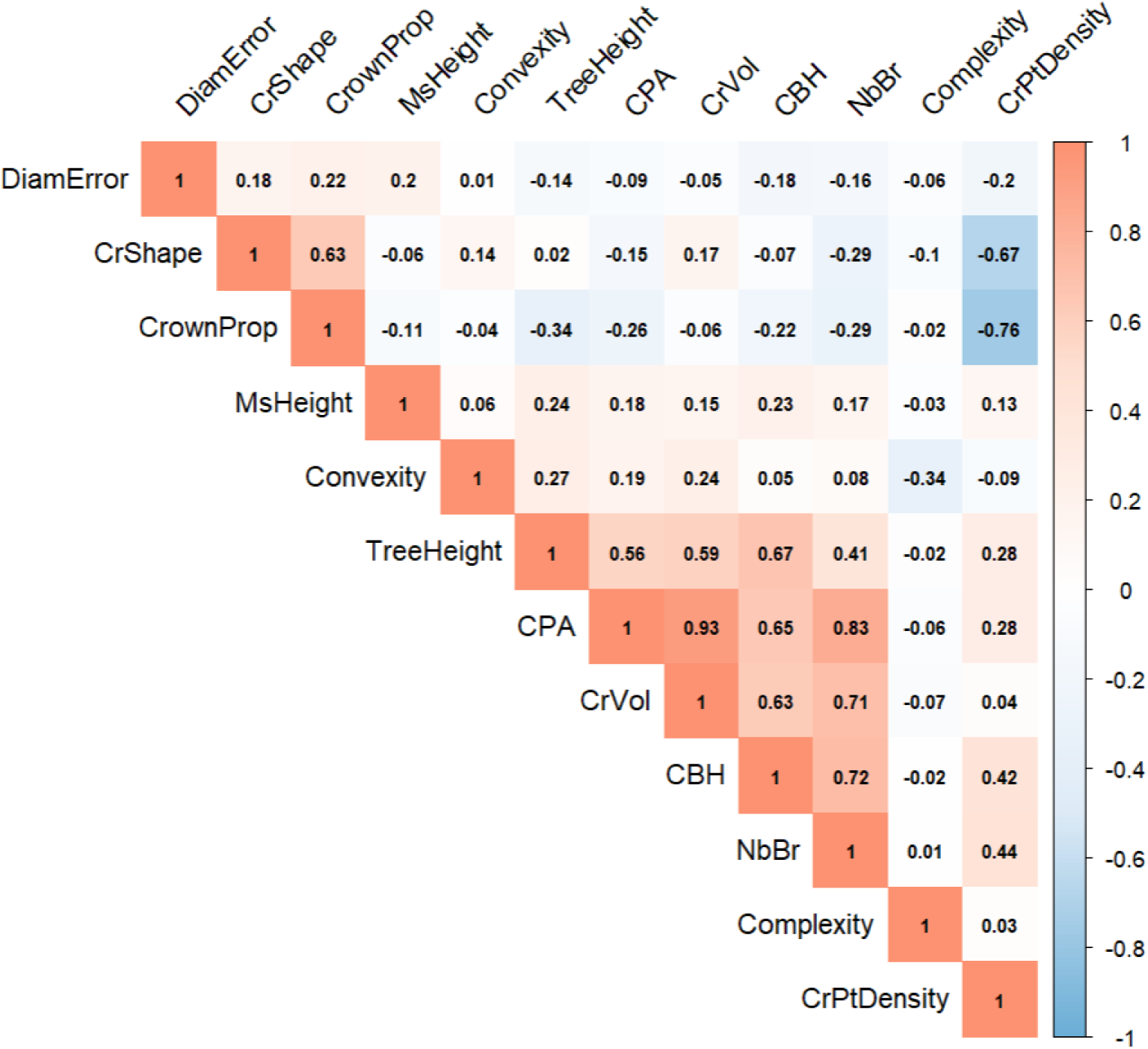
Correlation heat map between tree structural variables and diameter error, computed on the 7,218 pooled diameter measurements. Colours indicate the sign (red: positive, blue: negative) and the shade the strength of the Pearson correlation coefficient. DiamError: diameter error; MsHeight: height of the diameter measurement; CBH: circumference at breast height; NbBr: number of large branches; CPA: crown projected area; CrVol: crown volume; CrShape: crown shape; CrownProp: crown proportion; CrPtDensity: crown point density; Convexity and Complexity: stem convexity and complexity at breast height.

#### Tree level estimation (height and volume)

As for the diameter estimations, the principal components of the height and volume analysis were related to tree size, crown architecture and stem complexity, and errors were not grouped with these gradients (figure A4).

MLS-height errors were moderately correlated with stem complexity (r = −0.43), and weakly correlated with stem convexity (r = 0.17) and CBH (r = 0.15) (Figure 6). In contrast, the Vertex-height error was moderately correlated with more variables. The highest correlations were observed for CrVol (r = 0.26), CPA (r = 0.25), CBH (r = 0.25), NbBr (r = 0.23), CrownProp (r = −0.2), and CrPtDensity (r = 0.2).

**Figure 6:**
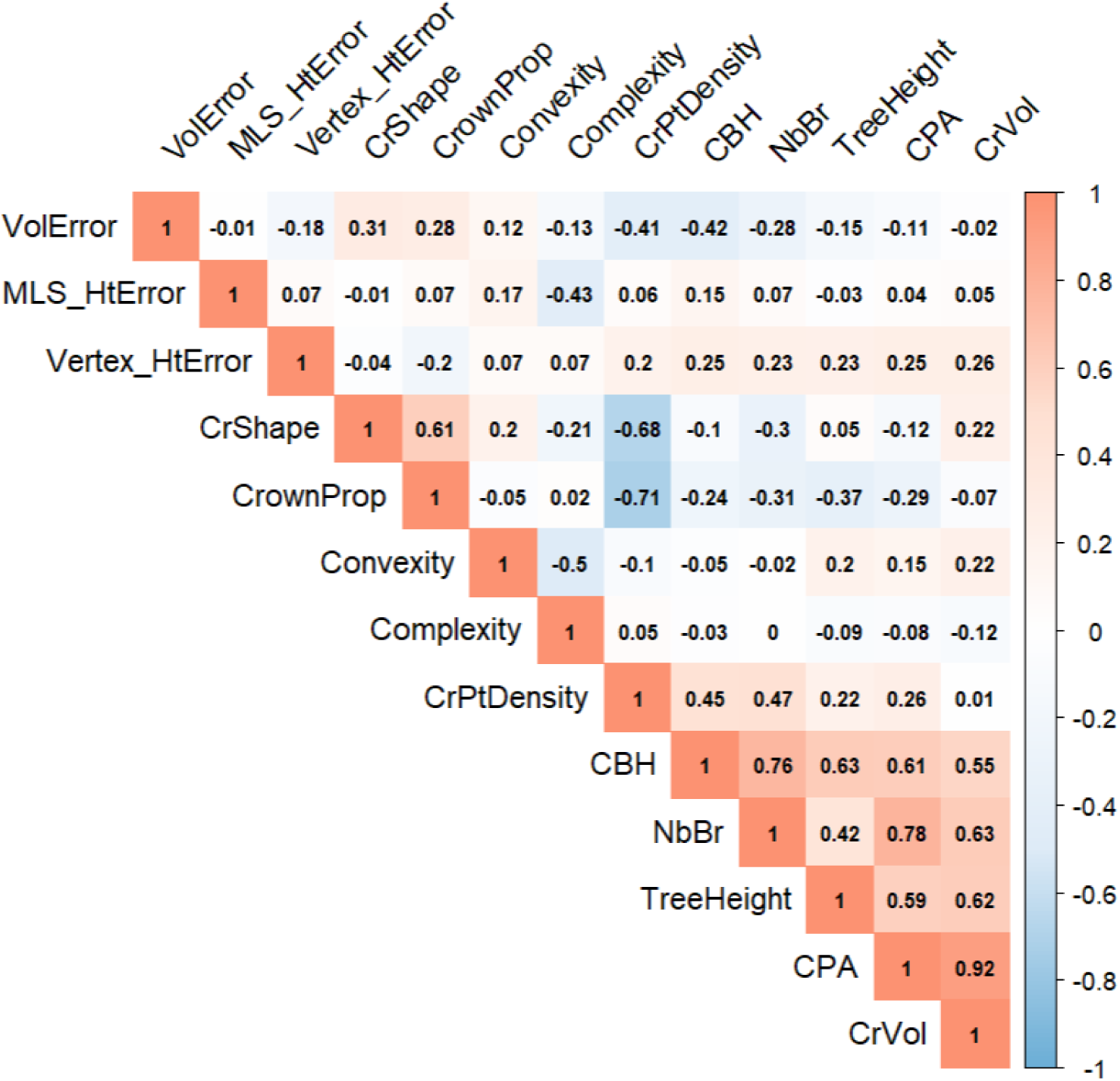
Correlation heat map between tree structural variables and the errors in tree height and stem volume, computed on 176 trees. Colours indicate the sign (red: positive, blue: negative) and the shade the strength of the Pearson correlation coefficient. MLS_HtError: height error of mobile laser scanning; Vertex_HtError: height error of the Vertex IV clinometer; VolError: error of the taper-based stem volume. Errors are estimates minus destructive reference measurements CBH: circumference at breast height; NbBr: number of large branches; CPA: crown projected area; CrVol: crown volume; CrShape: crown shape; CrownProp: crown proportion; CrPtDensity: crown point density; Convexity and Complexity: stem convexity and complexity at breast heigh

The volume error was moderately correlated with (CBH; r = −0.42), CrPtDensity (r = −0.41), CrShape (r = 0.3), as well as with CrownProp (r = 0.28). Larger trees with denser crowns generally exhibited lower volume estimation errors, whereas trees with more slender crown shapes tended to display greater measurement bias.

These correlations and principal components were computed on pooled observations and do not account for the nesting of measurements within trees, plots and species. They are therefore only indicative. The mixed-effects models below test the effect of tree shape while accounting for this structure.

### 3.3 Mixed-effects models of measurement errors

Final mixed-effects models are summarized in Table 8. Fixed effects alone explained 9.3–23.1% of the variance in measurement errors (marginal R^2^). Adding the random effects raised the explained variance to 29.4–39.3% (conditional R^2^), so more than 60% of the variance remained unexplained in every model.

**Table 8:**
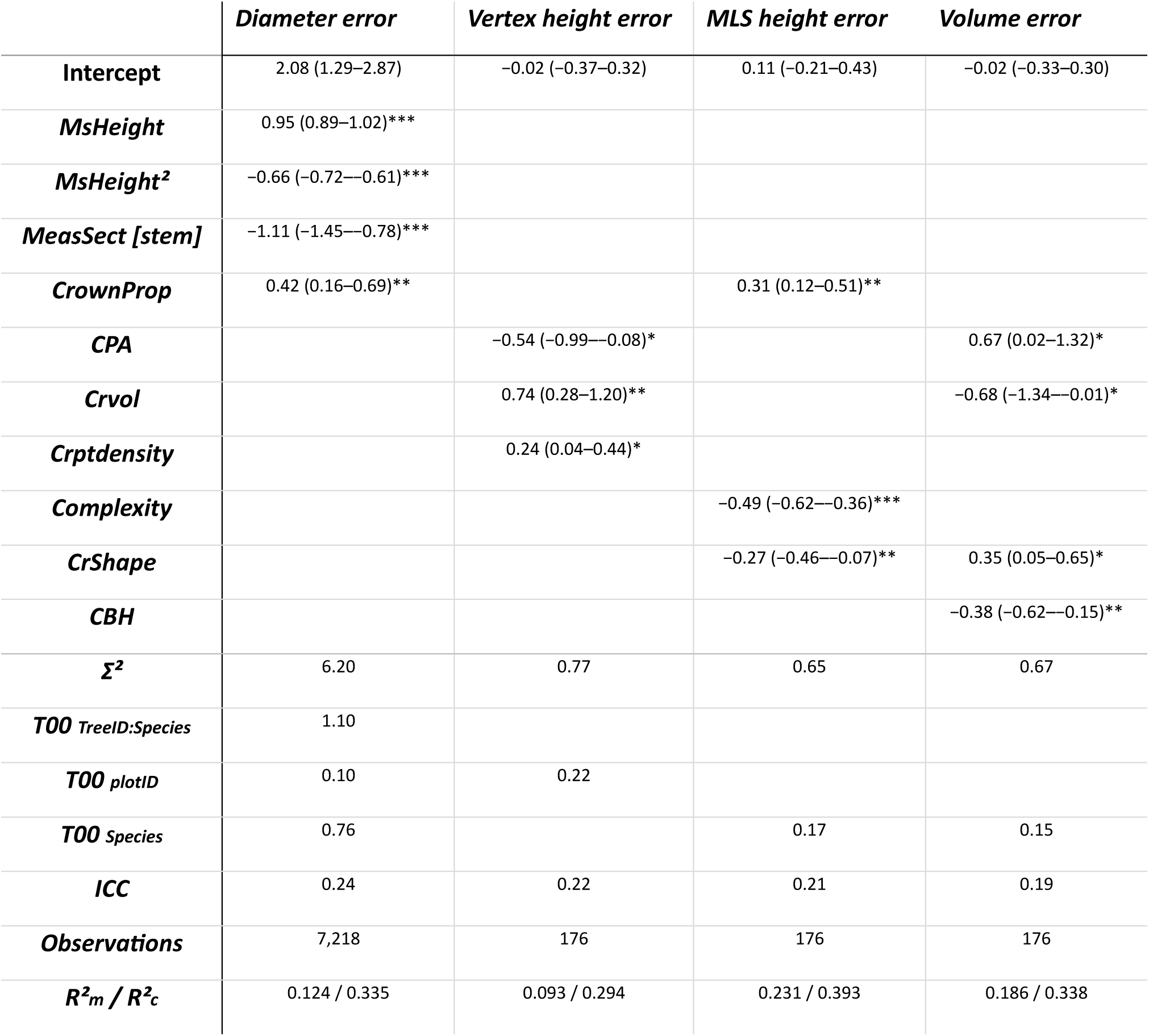
Linear mixed-effects models of the measurement errors. Estimates are standardized. The S5% confidence intervals is displayed in brackets. p: p-value; σ^2^: residual variance; τ00: random-intercept variance; ICC: intraclass correlation coefficient; R^2^m and R^2^c: marginal and conditional R^2^. *p < 0.05, ** p < 0.01, *** p < 0.001

The diameter error model retained the height of measurement (linear and quadratic terms, both p < 0.001), the measurement section and the crown proportion (CrownProp) as fixed effects (Table 8). Diameter error varied non-linearly with height, consistent with the underestimation near the base and overestimation higher along the stem reported in section 3.1.1. Errors were 1.11 cm lower in the stem section than in the crown section (p < 0.001), and increased with crown proportion (0.42, p = 0.002). Tree identity, species and plot were retained as random intercepts. Fixed effects explained 12.4% of the variance, and the full model 33.5%.

The Vertex IV height error was related to crown variables only: it decreased with crown projected area (p = 0.022) and increased with crown volume (p = 0.002) and crown point density (p = 0.020). Plot was the only random effect retained (ICC = 0.22). The MLS height error decreased with stem complexity (p < 0.001) and crown shape (p = 0.007) and increased with crown proportion (p = 0.002). For this model, species was retained as a random effect, and plot ID was not (ICC = 0.21). Fixed effects explained 9.3% (Vertex IV) and 23.1% (MLS) of the variance, and the full models 29.4% and 39.3%.

The taper-based volume error decreased with circumference at breast height (p = 0.001) and crown volume (p = 0.047) (Leading to more underestimation), and increased with crown projected area (p = 0.042) and crown shape (p = 0.021). Species was retained as a random effect and plot was not (ICC = 0.19). Fixed effects explained 18.6% of the variance and the full model 33.8%.

## 4. Discussion

### 4.1 Accuracy of mobile laser scanning

#### Diameters

MLS estimated stem diameters with a low overall bias, but the error depended strongly on the position along the stem. Diameters were underestimated under 3m, overestimated in the middle part of the stem, and became markedly less precise above about 15 m (Table 5, Figure 3). The same pattern was reported by Demol et al. (2021), Vandendaele et al. (2022) and Stovall et al. (2023), who also observed a loss of precision above 11.5 m. This is most likely due to i) the growing distance between the scanner and the stem, which widens the laser footprint through beam divergence and lowers point density, ii) occlusion by branches and foliage, which prevents the correct identification of stem points and prevent the diameter algorithm to converge, and iii) the movement of the upper stem in the wind, which makes the point cloud noisier.

The nature of the error also differed between species groups. Diameters were overestimated for softwood and almost unbiased for hardwood (Table 5), while the dispersion of the errors was similar. The difference therefore reflects a systematic overestimation in softwood rather than a lower precision. Several mechanisms could explain this. Needles and persistent whorled branches produce points close to the stem surface that are difficult to separate from true stem points, which increases the fitted diameter (Stovall et al., 2017). Bark roughness and stem deformation at branch attachments, which are more pronounced in conifers, also lead circle fitting to larger overestimates for spruce than for birch (Pitkänen et al., 2019). Finally, softwood species such as spruce and Douglas fir have lower crown bases than hardwood, so more of their diameters are measured within the crown, where branching leads to overestimation.

#### Height

MLS and the clinometer had similar overall precision, but their bias and relative performance depended on the species group (Table 6). In hardwood, MLS was more precise than the clinometer: its errors typically stayed below 2.5 m, whereas clinometer errors repeatedly reached 5 m. This agrees with studies reporting large clinometer uncertainty for hardwood trees, with maximum errors above 7 m for beech and a tendency to overestimate height (Stereńczak et al., 2019; Jurjević et al., 2020). In hardwood, the apex is difficult to identify visually from the ground within a complex, occluded crown, whereas MLS records the three-dimensional crown structure and can locate the highest points even when the apex is not identifiable from below. In softwood, both instruments underestimated height, and the clinometer was slightly more precise. The MLS underestimation was smaller than the one reported by Chiappini et al. (2022) for pine (up to 7.77 m), which may reflect differences in stand structure and tree architecture, or improvements in scanner range and performance. Overall, the advantage of MLS over clinometer measurements is context-dependent and varies with species group and crown structure.

#### Stem volume

The taper-based method gave nearly unbiased stem volumes and the best overall precision, with performance differing between species groups (Table 7). Volumes were slightly underestimated for hardwood and overestimated for softwood. This is consistent with the diameter results, since the systematic overestimation of softwood diameters propagates into volume. The relative error also depended on tree size: the taper method estimated smaller trees more accurately, while the relative error was smallest for the largest trees. These results are within the range reported in the literature for MLS (relative errors from −4.1% to 14.55% and RMSE% from 8.9% to 21.82%; Holvoet et al., 2025a; Chiappini et al., 2022; Vandendaele et al., 2022).

QSM volumes were systematically underestimated, and the underestimation increased with tree volume. It is probably partly related to the use of MLS instead of TLS, for which QSM were originally designed and validated (Raumonen et al., 2013): MLS point clouds are noisier and less homogeneous, with points both outside and inside the true stem surface, which disturbs cylinder fitting. A similar bias was reported by Vandendaele et al. (2022), who suggested applying a correction factor. A correction factor of 1.24, empirically tested, removed most of the bias for both species groups and gave errors comparable to the taper method, with the corrected QSM performing better for larger softwood trees. However, this factor is specific to our scanner and processing protocol and may not transfer to other equipment. It also had to be calibrated on our destructive reference data, which will often be unavailable in practice. Any change of scanner or protocol would therefore require a new calibration against reference measurements.

### 4.2 Influence of Tree Shape and architecture on MLS measurement error

#### Tree shape is not the main source of error

In the mixed-effects models, tree-level fixed effects explained only 9.3 to 23.1% of the variance in measurement errors, and 29.4 to 39.3% once random effects were included (Table 8). Correlations were weak to moderate, with |r| < 0.22 for diameter errors and |r| < 0.46 for height and volume errors, and the principal components did not group the errors with the main axes of structural variation (figure A4). More than 60% of the variance therefore remained unexplained in every model. This agrees with Panagiotidis et al. (2021), who found that the accuracy of laser scanning diameters depends mainly on acquisition factors (scan mode, scanner position, point density, processing method) and on local conditions such as wind and occlusion.

#### Diameters

Diameter error depended mainly on the position along the stem. The non-linear effect of measurement height matches the pattern described in section 4.1. To that is added the effect of the measurement section is added: diameters measured in the crown were on average 1.11 cm larger than those measured on the stem. This is consistent with the stronger occlusion in the crown leading diameter algorithm to fail to converge, and with a more irregular stem section where branches insert (Stovall et al., 2023). Crown proportion had a smaller positive effect on diameter error (+0.42 cm per standard deviation). This effect should be considered with caution; Trees with proportionally larger crowns have more diameters measured within the crown, so crown proportion and measurement section are not entirely independent. Crown proportion also differs between softwood and hardwood, since softwood trees have lower crown bases. Overall, diameter errors seem governed by local point cloud characteristics (crown or stem measurement, distance to the scanner, occlusion, noise) rather than by global tree attributes, in agreement with Panagiotidis et al. (2021), who found that laser scanning diameters depend mainly on acquisition conditions and processing rather than on tree structure.

#### Height errors

Vertex IV and MLS height errors were related to different variables, which suggests that the two methods are limited by different factors. For the clinometer, the error increased with crown volume and crown point density, which reflects the difficulty of locating the apex visually when it is hidden within a large, dense crown (Stereńczak et al., 2019; Larjavaara and Muller-Landau, 2013). Plot was the only random effect retained for the Vertex model, showing part of the error depend on site-specific conditions such as tree density or slope among others. For MLS, the error was related to crown shape, crown proportion and stem complexity, and the random effect was species. Narrow, elongated crowns, typical of softwood, increase the number of branch and foliage returns that the laser must cross before reaching the stem apex, which reduces the chance of capturing it. This interpretation is consistent with the lower MLS height performance on softwood (section 4.1), but it should be tested by mapping occlusion within the crown directly (Kükenbrink et al., 2025). The negative effect of stem complexity could reflect species-specific bark structure, such as the deeply furrowed bark of locust, or tree age, with older trees having more deeply carved bark, but our data cannot confirm it.

#### Volume

The taper-based volume error decreased with tree size (CBH) and crown volume, and increased with crown projected area and crown shape. Larger trees tended to have smaller relative errors, but a larger absolute underestimation. The positive effect of crown shape indicates that narrow, elongated crowns were associated with overestimation, in line with the positive softwood bias reported in section 4.1. Crown volume, crown projected area and crown shape are mathematically related (crown shape is crown volume divided by crown projected area), so their coefficients should not be read as three separate effects but should be considered together. Species was retained as a random effect, which points to species-specific traits not captured by our descriptors, such as bark texture, affecting the volume estimated from point clouds (Xie et al., 2025).

#### Species, site and unexplained variance

Species was retained as a random effect for the MLS height and volume errors, and plot for the diameter and clinometer height errors. Species-specific traits (branching pattern, foliage density, bark) and site-specific conditions therefore influence errors beyond the descriptors we tested. The large unexplained variance probably comes mostly from factors not tested here: acquisition conditions (trajectory, distance to the scanner), environmental conditions (stand density, slope, understory, phenology) and processing steps (segmentation, denoising, stem fitting). Tree shape did not allow errors to be predicted for an individual tree, but it shows where MLS is more uncertain: within the crown, for narrow-crowned softwood trees, and for the smallest trees.

### 4.3 Limitations and perspective

#### Limitations

The main limitation of this study is that it tested only individual tree descriptors as predictors of MLS measurement errors. Environmental conditions and acquisition parameters were not tested. Stand density, slope, understory vegetation, leaf phenology or the distance to the scanning trajectory are all plausible drivers of point cloud quality, and acquisition variables such as walking pattern, walking speed, loop closure and beam divergence were not accounted for. Their effect cannot be excluded, and it may interact with tree form and stand structure. Vandendaele et al. (2024) showed, for example, that the acquisition pattern had a limited effect on height, diameter and crown attributes but influenced volume estimates, with denser acquisitions increasing noise-related overestimation. This suggests that the relationship between acquisition design and measurement accuracy is nuanced and variable-dependent, and may interact in complex ways with stand structure and tree form. A second limitation comes from the nested statistical structure. Finally, the data were collected in leaf-off conditions in a single region (Wallonia), so the results may not transfer to other forest types or seasons.

#### Contribution and perspectives

Despite these limitations, the 176 destructively sampled trees, covering eight species, a wide range of sizes and varied tree shapes, provide a comprehensive reference for MLS validations. An operational use of MLS in national and regional forest inventories (Holvoet et al., 2025a; Kükenbrink et al., 2025) requires not only acceptable mean errors, but also knowing under which conditions errors increase. This study contributes by showing that tree shape alone is a weak predictor of error, which directs the search for error sources toward acquisition and environmental conditions. Future studies should therefore test tree characteristics, environmental variables and scanning specifications simultaneously, in a wider range of forest types, stand structures and regions. This would allow models predicting measurement errors, which could flag unreliable measurements during post-processing and help to optimize acquisition protocols and to make reporting more robust.

## 5. Conclusion

Using 176 destructively sampled trees of eight species, we evaluated the accuracy of mobile laser scanning (MLS) for stem diameters, total tree height and merchantable stem volume. Diameters were estimated with a low overall bias (+0.5 cm), but precision decreased with height along the stem, particularly above 15 m, and diameters were overestimated for softwood. Tree height was estimated with a precision similar to that of a clinometer, with a clear advantage for hardwood and a slight disadvantage for softwood. A taper-based reconstruction gave nearly unbiased stem volumes (RMSE% of 15.7), whereas volumes from quantitative structure models were systematically underestimated (−21.2%).

Tree shape influenced measurement errors but explained only a limited part of them. In mixed-effects models, tree architecture variables explained 9.3 to 23.1% of the variance in errors, and 29.4 to 39.3% once species and plot ID were included. Errors were larger within the crown, for narrow-crowned softwood trees, and differed with tree size, whereas clinometer height errors were related to crown volume and density. None of the diameter, height or volume errors could be predicted from tree shape alone, which indicates that tree shape is not the sole source of MLS measurement uncertainty.

Most of the error therefore seems to originate from other sources, such as scanning conditions, stand and site characteristics and processing steps. Before MLS is used in large-scale forest inventories, future studies should vary these factors together with tree characteristics, while mapping occlusion within the crown directly.

All in all, this study moved the use of MLS closer to operationalization by exploring the impact of individual tree characteristics on measurement error and showing that MLS measurements are relatively robust across a wide range of shape and size of trees. It also showed that a bigger focus will need to be put on environmental conditions and scanning protocol in order to fully understand the source of MLS measurement error.

## 6. Data availability statement

The dataset presented in this paper including the individual tree scans of the 176 trees, are available in open access (CC BY 4.0 license) on the ULiège Dataverse server at the following DOI: https://doi.org/10.58119/ULG/HI1CLM.

The script used to produce the result presented in this article are available at https://github.com/HolvJ/MLSVolumeEstimations

## 7. Acknowledgement

This research was founded by the Interreg Grande Région project W.A.V.E. (Wood Added Value Enabler, https://wave-gr.eu/en/). The authors would like to thank Florentin Reginster, Alan Borremans, Boris Lemaigre, Tom Mortelmans, Charlotte Longrée, Pauline Cubelier, Adèle Philipo, Gladis Werenne, Maxence Wagnon, and Marie-Pierre Tasseroul for the necessary help during the measurement on the felled trees. We also thank Mr. Eric de Rese for allowing us to scan and measure the felled trees in the Seraing Arboretum, Mr. Camille Deleau for the trees near Verlée, and Mr. Philippe Cornet for the spruces and Douglas firs.

## Appendix

**Table A1:** Complete list of the variables used in the analyses, with their origin and the way they were obtained. Abbreviations are defined in Table 3.

| **Variable** | **Abreviation** | **Description** | **Origin** | **Obtained by** |
| --- | --- | --- | --- | --- |
| **Number of large branches** | *NbBr* | *The number of branches of the tree with initial circumference above 20 cm.* | *Manual measurement* | *Counted on the felled trees.* |
| **Circumference at breast height** | *CBH* | *Circumference of stem at 1.30m above ground in cm.* | *Manual measurement* | *Using a measuring tape.* |
| **Tree total height** | *TreeHeight* | *The manually measured tree height on felled tree* | *Manual measurement* | *Measuring the distance between the base of the trunk and the top of the stem* |
| **Stem merchantable volume** | *StemVolume* | *The manually measured stem merchantable volume* | *Manual measurement* | *Summing the volume of each stem section measure on the field* |
| **Vertex stem height estimate** | *Vertex_Height* | *The height of standing trees measured by a Vertex IV* | *Manual measurement* | *Using the Vertex IV* |
| **MLS stem diameter estimate** | *MLS_Diameter* | *The MLS estimates of successive stem diameters* | *Point Cloud* | *Applying the ITSMe circle fitting algorithm on the point cloud on successive 10cm slices* |
| **MLS stem height estimate** | *MLS_Height* | *The MLS estimates of tree height* | *Point Cloud* | *Calculating the Z axis difference between the highest and lowest point of the individual tree point clouds* |
| **MLS stem taper merchantable volume estimate** | *MLS_Volume* | *The MLS estimation of stem merchantable volume acquired by integrating the individual stem taper equations* | *Point Cloud* | *Fitting a taper equation on the diameter measurements and integrating this equation from the base of the trunk up to the MLS_Height.* |
| **Diameter measurement height** | *MsHeight* | *The height of the diameter measurement.* | *Point Cloud* | *Calculating the Z axis difference between the base of the point cloud and the diameter measurement.* |
| **Stem convexity at breast height** | *Convexity* | *Index defined as the ratio between the area of the spline fitted on the slice of the point cloud and the convex hull fitted on the same slice.* | *Point Cloud* | *Calculated using the method presented in Bauwens et al., (2021).* |
| **Stem complexity at breast height** | *Complexity* | *Index describing the complexity of the stem at breast height.* | *Point Cloud* | *Dividing the circumference of the circle fitted at breast height by the perimeter of the spline fitted at breast height previously acquired.* |
| **Crown projected area** | *CPA* | *Surface of the convex hull formed around the crown point projected on a horizontal plane in m<sup>2</sup>.* | *Point Cloud* | *Using ITSMe R package.* |
| **Crown volume** | *CrVol* | *Volume of the alpha shape build around the crown points in m<sup>3</sup>.* | *Point Cloud* | *Using ITSMe R package.* |
| **Crown/stem ratio** | *CrownRatio* | *The ratio between total tree height and crown base height.* | *Point Cloud* | *Using ITSMe R package.* |
| **Crown point density** | *CrPtDensity* | *Index describing the point density in the space occupied by the crown.* | *Point Cloud* | *Dividing the number of crown points by the crown volume* |
| **Crown shape parameter** | *CrShape* | *The ratio between crown volume and crown projected area* | *Point Cloud* | *Dividing the Crown volume by the Crown projected area* |
| **MLS estimates diameter error** | *DiamError* | *The error of measurement of MLS diameter estimates compared with reference destructive measurement* | *Calculated from two measurements* | *Subtracting the manual diameter measurement from the MLS estimated diameter* |
| **MLS height estimate error** | MLS_HtError | The error of measurement of MLS height estimates compared to reference destructive measurement | Calculated from two measurements | Subtracting the destructive height measurement to the MLS estimated height |
| **Vertex height estimate error** | Vertex_HtError | The error of measurement of Vertex height estimates compared to reference destructive measurement | Calculated from two measurements | Subtracting the destructive height measurement to the Vertex estimated height |
| **MLS taper stem merchantable error** | VolError | The error of measurement of MLS taper merchantable volume estimates compared to reference destructive measurement | Calculated from two measurements | Subtracting the destructive merchantable volume measurement to the MLS taper estimated merchantable volume |

**Figure A1:**
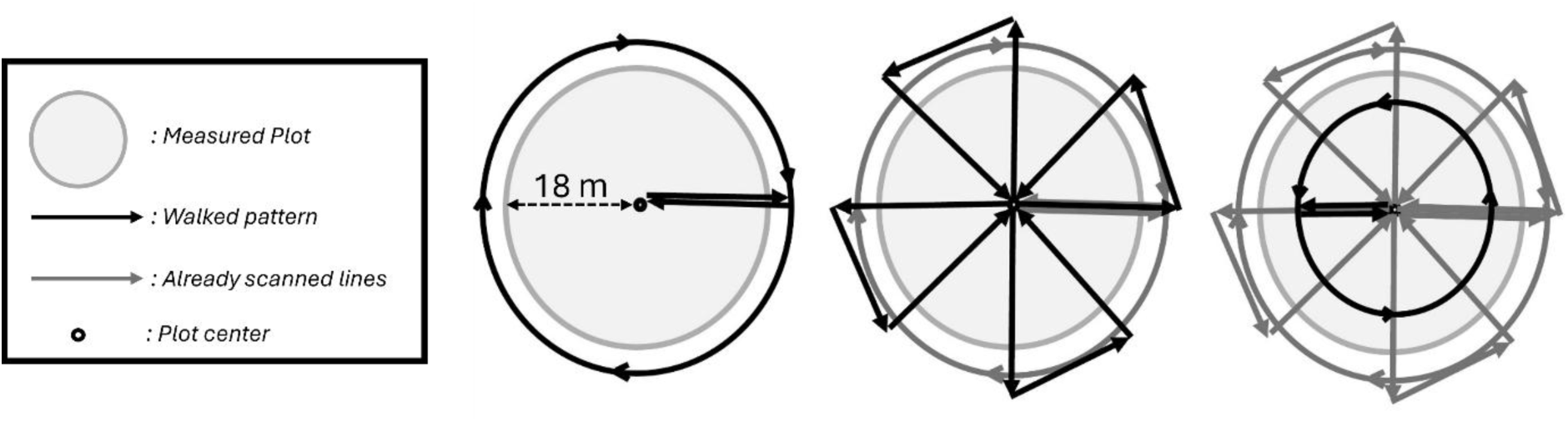
Walking pattern followed during the mobile laser scanning of plots. Black lines indicate, from left to right, the path taken; grey line indicates the path already covered; black dot indicates both the centre of the plot and the start and end points of the scan; the lighter grey circle delimits the surveyed plot.

**Figure A2:**
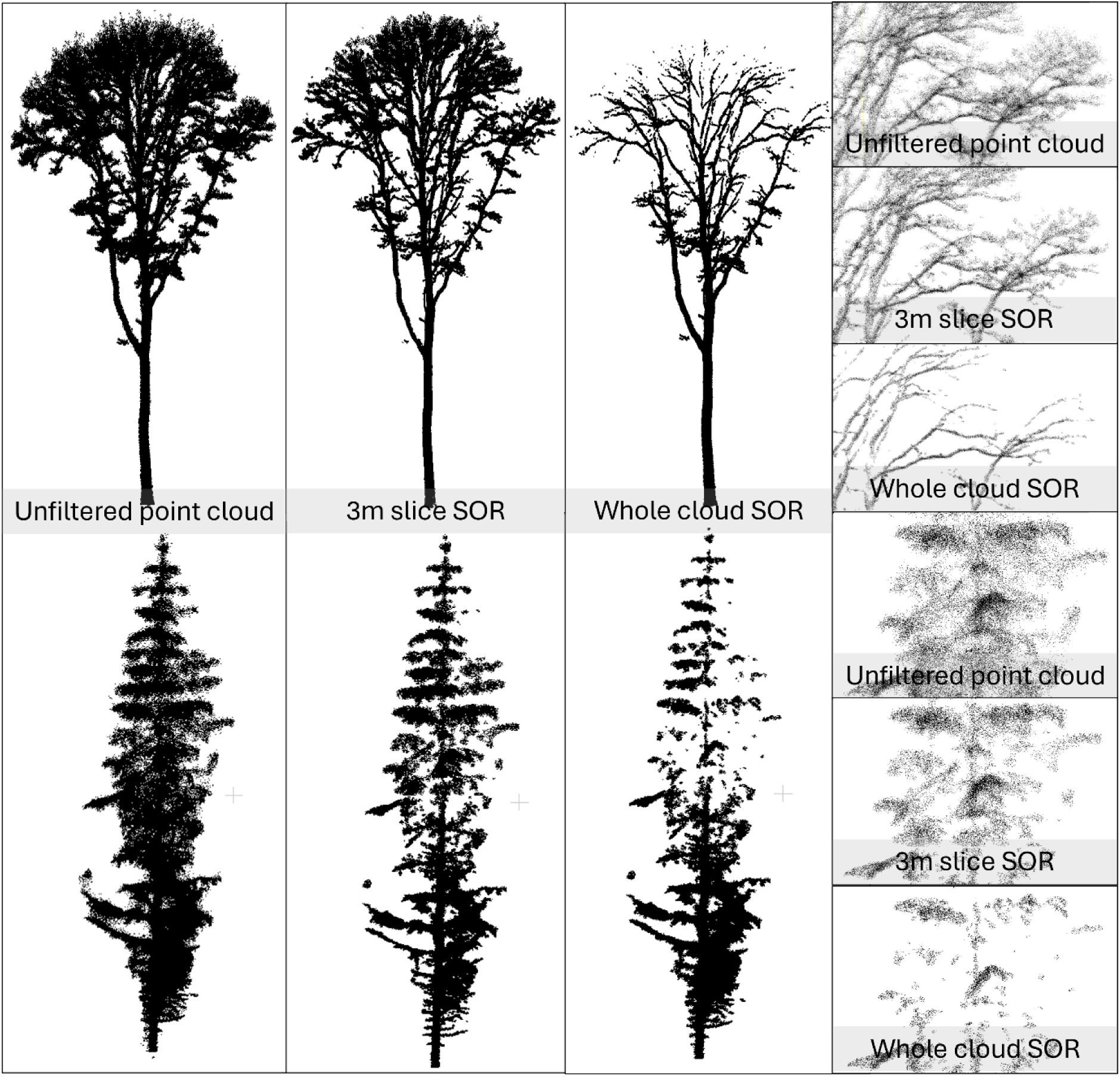
comparison of noise removal between statistical outlier removal applied to the whole tree point cloud and by 3 m height slices on a Spruce and Beech point cloud.

**Figure A3:**
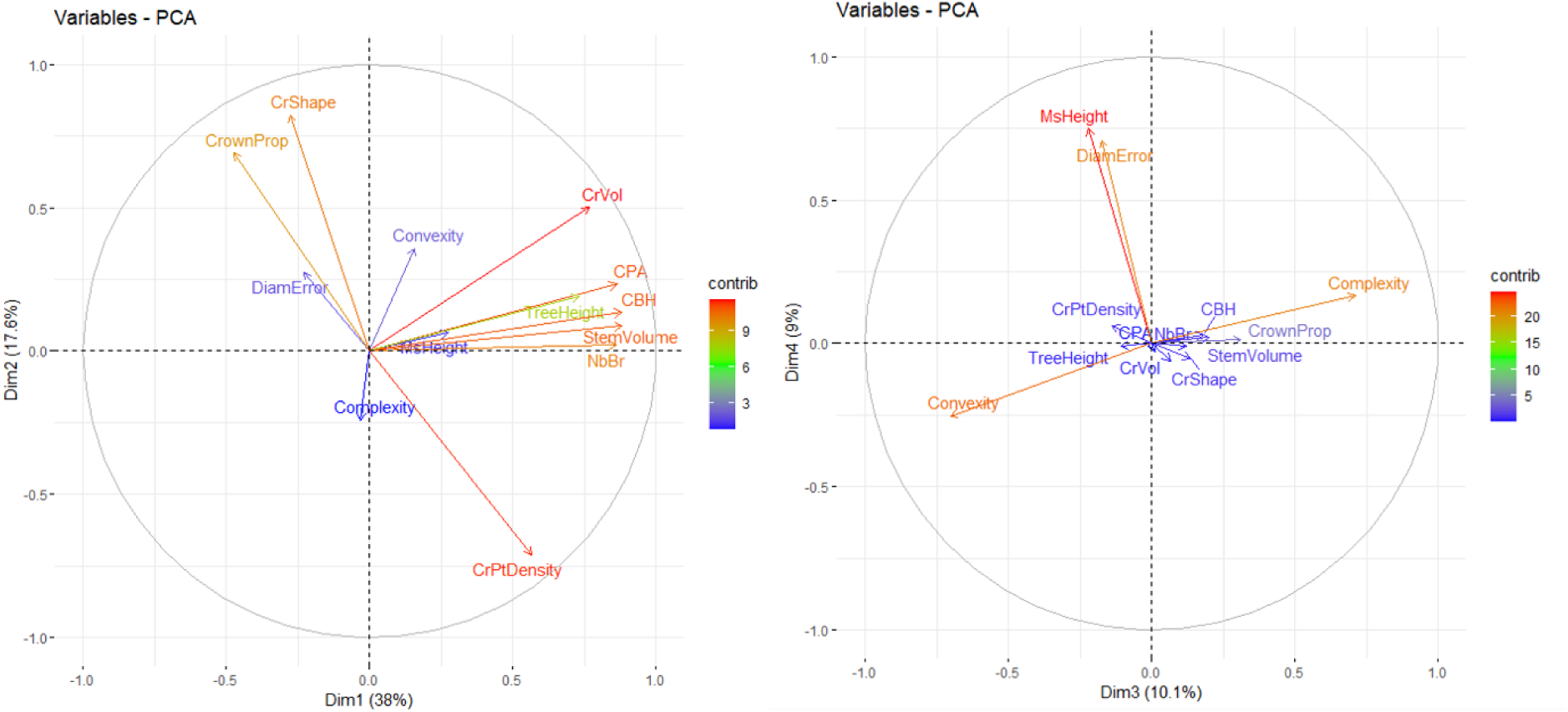
Result of the diameter estimation PCA build on the 7218 diameters measurements. The first two components of the PCA are used in the left plot, the third and fourth component in the right plot. The color and length of the axes correspond to their contribution to the PCA.

**Figure A4:**
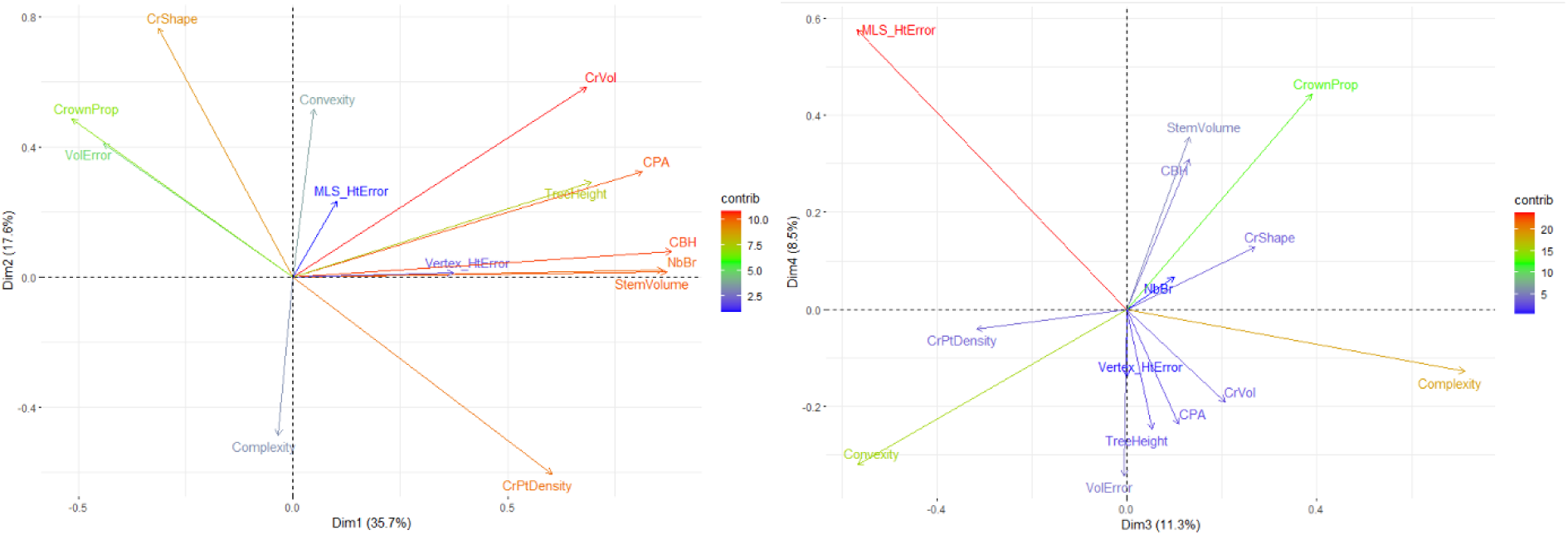
Result of the volume and height estimation PCA build on the 176 individual trees. The first two components of the PCA are used in the left plot, the third and fourth component in the right plot. The color and length of the axes correspond to their contribution to the PCA.

